# Peptide fibrils as vaccine: a proof of concept

**DOI:** 10.1101/2023.12.30.573725

**Authors:** Yana Zabrodskaya, Konstantin Sivak, Maria Sergeeva, Andrey Aleksandrov, Elena Kalinina, Alexandr Taraskin, Mikhail Eropkin, Elena Eropkina, Vladimir Egorov

**Author notes:** Corresponding author: Dr. Yana Zabrodskaya,;, Institute of Biomedical Systems and Biotechnology, Peter the Great Saint Petersburg, Polytechnic University, 29 Ulitsa, Polytechnicheskaya, St. Petersburg 194064, Russia.

## Abstract

The potential of amyloid-like fibrils formed by peptides as vaccine candidates was investigated using a fragment of the Ebola virus glycoprotein. Peptide in fibrillar form were found to induce an immune response to the full-length protein without causing cellular toxicity or significant changes in hematological studies. The ability of the studied peptide fragment to oligomerize and form amyloid-like fibrils and intermediates suggests potential implications for the virus’s mechanisms of action on cells, particularly those of the immune system. Additionally, if native GP2 epitopes are retained in the peptide fibrils, they may serve as effective immunization agents due to their autoadjuvant properties; however, it is important to consider the possibility of cross-reactivity with human proteins. These findings provide valuable insights into the potential use of amyloid-forming peptide as vaccine candidates and highlight the need for further research into their immunogenic and adjuvant properties.

## 1. Introduction

The availability of platforms that enable the rapid development of a vaccine to protect against pathogens causing local outbreaks is an urgent task, given the ongoing threats of epidemics and pandemics caused by new zoonotic infections. One such pathogen is the Ebola virus [1]. The experience gained from the pandemic caused by SARS-Cov2 has demonstrated that having platforms for vaccine development allows for a quick response to emerging challenges and facilitates the production of a vaccine against a completely new pathogen [2,3]. A flexible and rapidly deployable platform is the peptide vaccine platform [4].

However, the use of peptide vaccines presents several challenges, including low immunogenicity and the need to use adjuvants or create chimeric proteins to adapt the antigen structure for recognition by the human immune system [5,6]. In this study, we propose an approach to enhance the immunogenicity of peptide vaccines by utilizing the self-organization of peptides into spatial structures necessary for triggering an immune response. One such approach involves the formation of a quaternary structure through amyloid-like self-organization. We hypothesize that the induction of fibrillogenesis is specific, and the presence of immunogenic components of the virus that have formed fibrils cannot induce fibril formation by human proteins in the absence of homology between the antigen and host proteins [7]. To test this hypothesis and develop a vaccine capable of inducing the formation of virus-neutralizing antibodies, it is necessary to select a surface antigen with the following properties: (1) it must be located in a critical region, the interaction of antibodies with which leads to virus neutralization; (2) it must be capable of forming amyloid-like fibrils.

In a previous study [5], a spatial epitope of the Ebola virus GP protein was identified through epitope screening, and antibodies targeting this epitope were found to have a neutralizing effect. Additionally, our previous research demonstrated that a component of this epitope, the non-glycosylated GP region, is capable of forming amyloid-like fibrils [8]. Briefly, we found that a peptide matching the primary structure of the viral GP protein fragment can exist in soluble and fibrillar forms, depending on the amino acid residue at the C-terminus. A peptide with a Q at the C-terminus formed fibrils, while a peptide with a G at the C-terminus did not. This system allowed us to test the concept of enhancing the immunogenicity of potential peptide vaccines through the formation of amyloid-like fibrils by antigens. Therefore, in this study, we aimed to determine (1) whether the peptide contained in fibrils would induce the formation of immunoglobulins G to whole GP better than the peptide in soluble form, and (2) whether the introduction of fibrils into a model animal would lead to acute toxicity.

## 2. Experimental

### 2.1 Peptides and protein

Peptides ^551^QNALVCGLRQ^560^ (G33) and QNALVCGLRG (**G31**) were synthetized by LTD “Verta” (purity > 98%). To prepare peptides’ solutions 1 mg of peptides were dissolved in 1 mL of PBS (phosphate buffered saline, Sigma). The recombinant eGP protein obtained from a eukaryotic system (CHO cells) was generously provided by the Research Institute of Highly Pure Biological Preparations.

### 2.2 Cell viability assay

Cell viability of MDCK or A549 cells (ATCC) was determined using MTT assay (MTT, 3-(4,5-Dimethyl-2-thiazolyl)-2,5-diphenyl-2H-tetrazolium bromide, Sigma) according to manufacturer’s instructions. For the test 8 two-fold dilutions of G31 and G33 peptides’ solutions were used (from 500 ug/ml to 3.9 ug/ml). Optical density at 490 nm was measured using CLARIOstar (BMG Labtech).

### 2.3 Immunization

All animal procedures were conducted in compliance with the regulations for the ethical treatment of animals as outlined by the European Convention for the Protection of Vertebrate Animals Used for Experiments and Other Scientific Purposes (Strasbourg, 1986). White mice of the balb/c line from Rappolovo, Russia, were utilized as model animals. Peptides were administered intraperitoneally in PBS at a dosage of 1 mg/kg. The study included 3 groups of animals: 5 mice in the control group were given a single injection of PBS, 5 mice received an injection of a G31 peptide solution in PBS at a dose of 1 mg/kg, and 5 mice were injected with a G33 peptide solution in PBS at a dose of 1 mg/kg. The weight of the animals was monitored for 7 days. Following this period, serum ELISA was performed to detect specific immunoglobulins to the recombinant protein eGP (2 mice per group), along with a clinical blood test and histological analysis of the lungs, spleen, intestines, kidneys, and liver (5 mice per group).

### 2.4 ELISA

The eGP protein of the Ebola virus was adsorbed onto the wells of a 96-well plate at a concentration of 1 μg/ml. The plate was then incubated at 4°C overnight. After washing three times with PBST (PBS containing 0.05% Tween-20) and blocking with 5% Blocking reagent (BioRad) in PBST, mouse blood sera at various dilutions were added to the wells of the plate and incubated for 2 hours at 37°C. Antibodies bound to the recombinant protein were detected using a GAM-HRP secondary antibody conjugate (BioRad). Subsequently, the ELISA results were determined using a peroxidase reaction. Absorbance at wavelengths of 450 and 620 nm was measured using a CLARIOstar multimodal reader (BMG Labtech).

### 2.5 Hematological studies

Blood was analyzed for red blood cell count, hemoglobin concentration, hematocrit, average hemoglobin content in a cell, average hemoglobin concentration in a cell, white blood cell count, leukocyte formula, and platelet count using an automatic hematology analyzer Abacus Junior Vet (Austria).

### 2.6 Histology

After planned euthanasia (decapitation under anesthesia), the animals underwent autopsy to assess the damaging effect on internal organs and tissues. The external state of the body and the chest and abdominal cavity with organs and tissues were thoroughly examined, and all deviations from the norm were documented.

Histological studies of animal organs from all groups involved formalin fixation and histoprocessing. The lungs, spleen, intestines, kidneys, and liver were subjected to histological examination. Tissues were fixed in 10% buffered formalin and processed using standard biopsy cassettes on a Histo-Tek VP1 automated histoprocessor, followed by embedding in a paraffin block in stainless steel embedding molds. 5-μm sections were cut from paraffin blocks and stained with hematoxylin and eosin. The histological preparations were then examined under a light microscope.

## 3. Results and discussion

### 3.1 Peptides

To investigate the potential use of peptides capable of forming amyloid-like fibrils as vaccine candidates, a peptide fragment of the Ebola virus GP2 protein (^**551**^**QNALVCGLRQ**^**560**^, ***G33***), known to form amyloid-like fibrils, was synthesized. Figure 1 illustrates the location of the G33 peptide in the GP2 protein structure. Additionally, to assess the impact of the amyloidogenic component on antibody production, a peptide analogue (**QNALVCGLRG, *G31***) was synthesized with the C-terminal Q replaced by G, which exhibits lower propensity for fibrillogenesis and forms compact low-molecular oligomers. [8].

**Figure 1.**
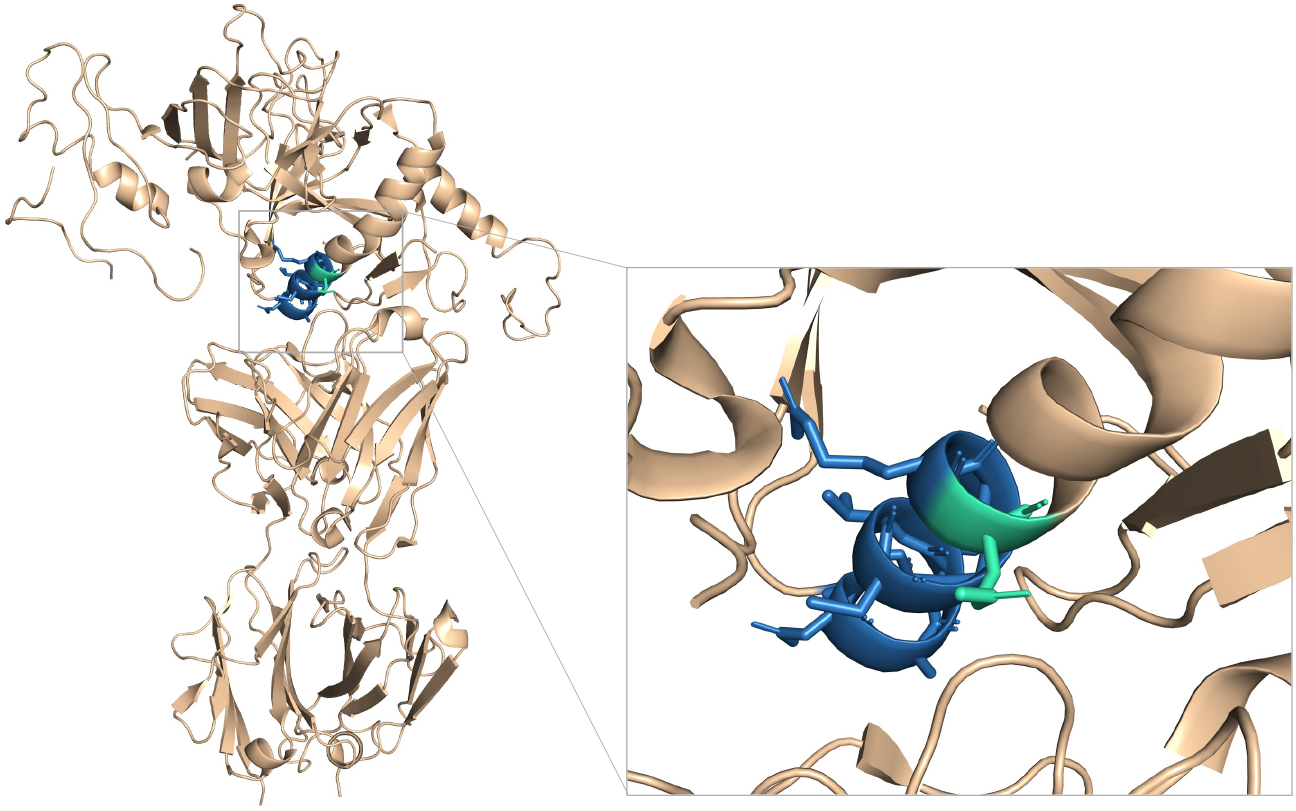
The location of the G33 peptide in the GP protein structure was determined using the crystal structure of Sudan Ebolavirus Glycoprotein (strain Boniface) 3VE0 for reference [9]

### 3.2 The toxicity of peptides G31 and G33

In order to test the hypothesis about the immunogenicity of G33 fibrils, initial experiments were conducted to assess the toxicity of the G31 and G33 peptides in cell cultures. The toxicity study of G31 and G33 peptides was performed on MDCK and A549 cell cultures, which are canine kidney epithelial cells and human lung carcinoma epithelial cells, respectively. The results of the MTT test indicated that these peptides were not toxic to the tested cell types up to the maximum concentration of 500 μg/ml (Figure 2).

**Figure 2.**
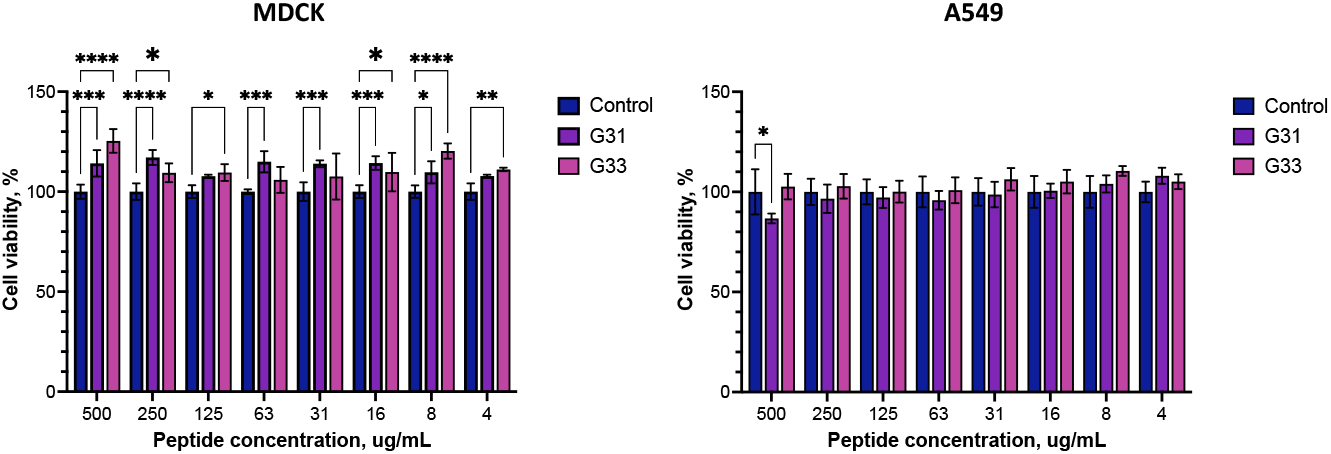
The results of the MTT test conducted on MDCK and A549 cells. ‘Control’: intact, ‘G31’ and ‘G33’: cells to which solutions of the G31 and G33 peptides were added. Statistical data processing was carried out using ANOVA adjusted for multiple comparisons (* – p < 0.05, ** – p < 0.01, *** – p < 0.001, **** – p < 0.0001)

The absence of toxicity in the cell model allowed for the initiation of an experiment to investigate the impact of introducing the studied peptides on animal models.

### 3.3 The immunogenicity of peptides G31 and G33

The immunization of balb/c mice was conducted using solutions of peptides G31 (in nonfibrillar form) and G33 (in fibrillar form) in PBS, with PBS utilized as a negative control. Observations of the body weight dynamics over 7 days did not reveal statistically significant differences between the groups. Following the 7-day period, serum ELISA was performed to detect specific immunoglobulins to the recombinant eGP protein, as well as a hematological studies and histological analysis of the lungs, spleen, intestines, kidneys, and liver.

ELISA demonstrated a increase in the titer of specific antibodies to the recombinant eGP protein in the sera of mice injected with the G33 peptide in comparison to the antibody titer in the sera of control animals and those injected with the G31 peptide (Figure 3).

**Figure 3.**
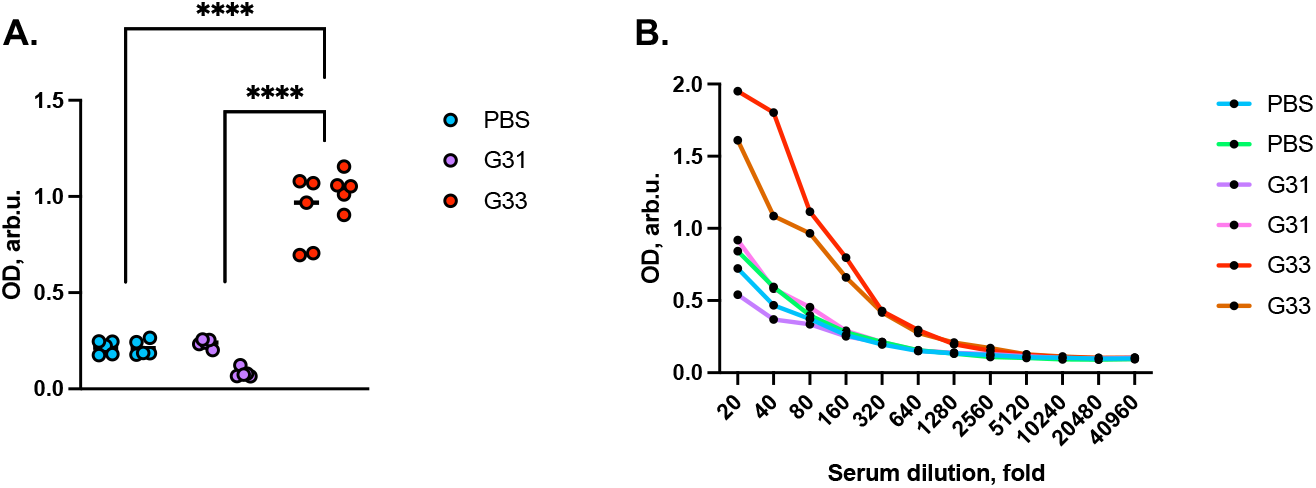
ELISA results (A) at a serum dilution of 1/100, (B) at various dilutions of blood sera from immunized mice. Statistical data processing was carried out using ANOVA adjusted for multiple comparisons (**** – p < 0.0001)

These results indicate that immunization with the G33 peptide is capable of stimulating the production of specific antibodies that recognize the recombinant protein eGP of the Ebola virus, highlighting the autoadjuvant properties of the peptide fibrils.

### 3.4 Evaluation of the effect of peptides on the mouse body

The introduction of the G31 peptide resulted in a suppression of lymphocytic and mononuclear components, along with a tendency for decreased granulocytes in the leukocyte formula. The number of leukocytes and mononuclear cells changed by more than twice against the background of leukopenia in the control group, but all values remained within the physiological norm for mice (Figure 4). However, there were no significant changes observed in red blood counts, and no signs of anemia were detected. Furthermore, there were no substantial differences observed in the platelet component of cellular hemostasis.

**Figure 4.**
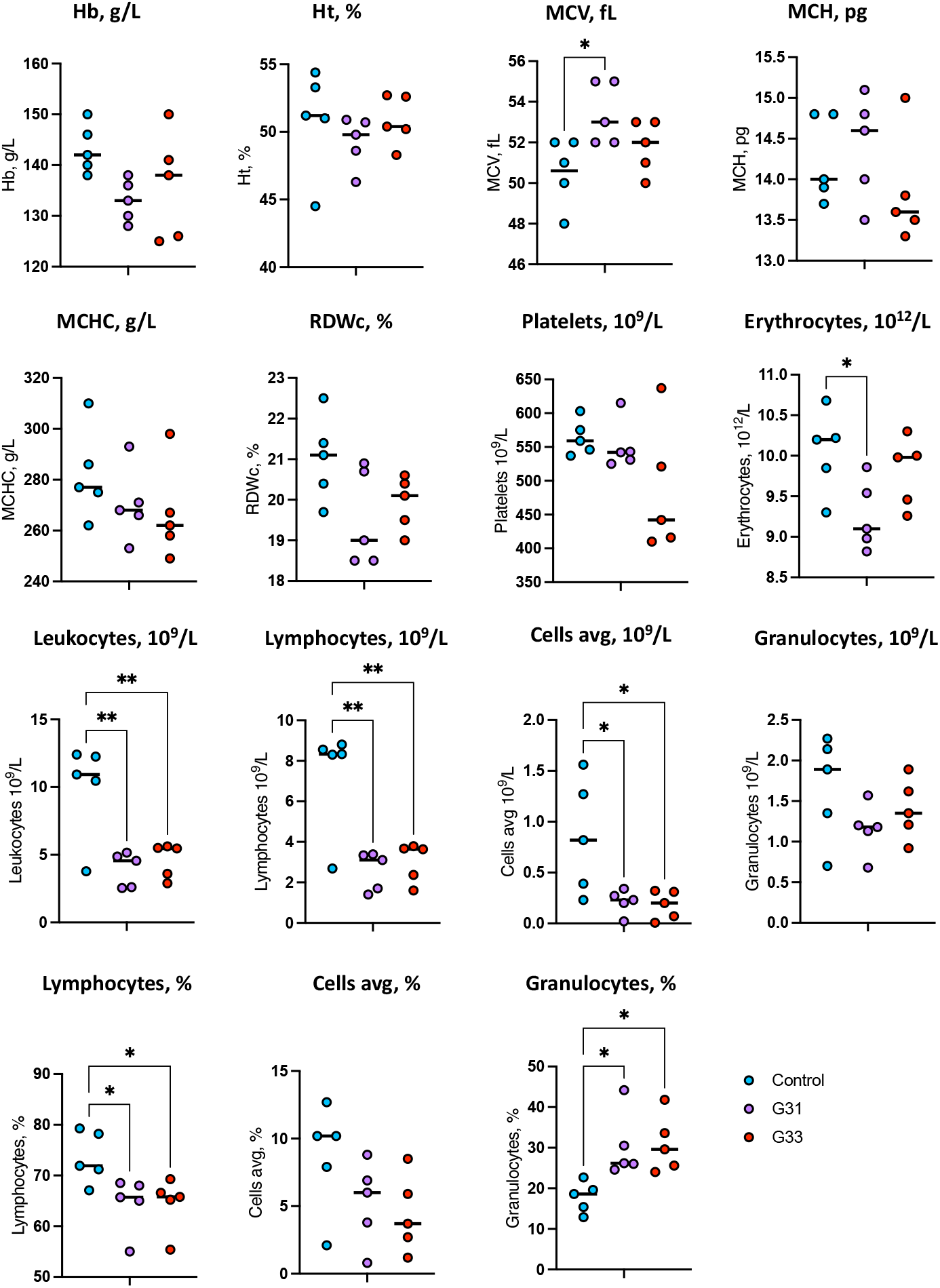
Results of hematological studies. ‘Control’ – immunization with PBS, G31 and G33 – immunization with solutions of the corresponding peptides. Statistical data processing was carried out using ANOVA adjusted for multiple comparisons (* – p < 0.05, ** – p < 0.01).

When examining the histological preparations of the liver and kidneys, interstitial infiltrates were identified (Figure 5). The structure of the remaining studied organs was within normal limits.

**Figure 5.**
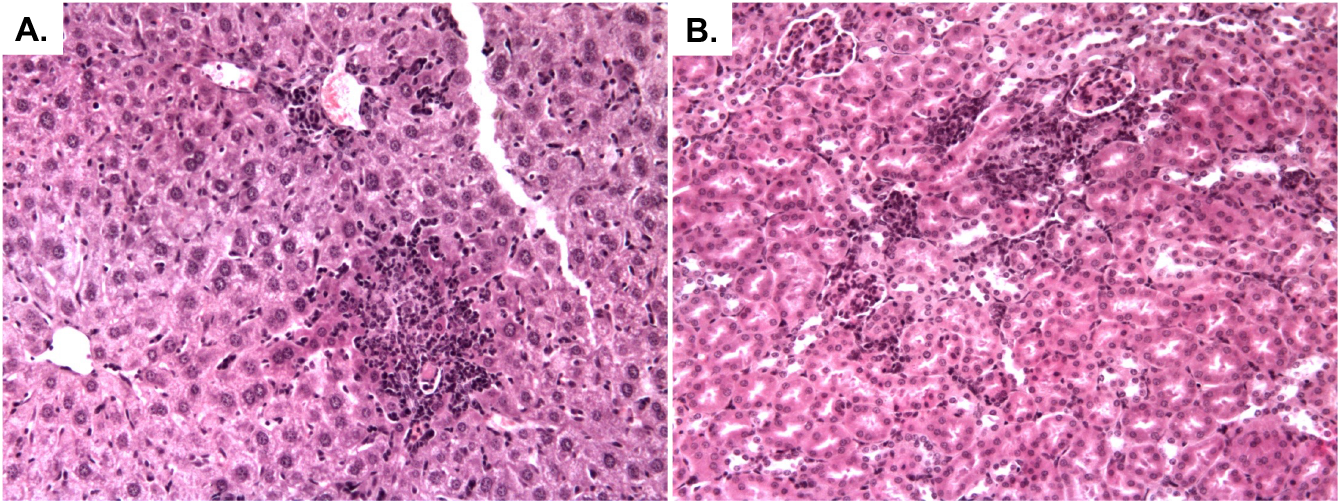
**(A)** Morphological structure of the mouse liver on the 7th day after administration of the G33 peptide. Increased cytosis and extensive foci of periportal infiltration. **(B)** Morphological structure of the mouse kidney on day 7 after administration of peptide G33. Foci of lymphoid-macrophage infiltrate around the glomeruli and renal tubules. Stained with Hematoxylin-eosin, magnification ×240.

In general, the pathological changes in organs were qualitatively similar when using both peptides, but quantitatively these changes were more pronounced in animals receiving the G33 peptide. The main changes were observed in the liver and kidneys. In the livers of animals receiving the G33 peptide, an increased number of Kupffer cells was noted, along with extensive areas of cellular infiltration primarily localized periportally. Hepatocytes appeared intact, with undisturbed structure, and no increased vacuolization was observed (Figure 5A).

A similar pattern was observed in the kidneys of the animals, where cells of the lymphoid-macrophage infiltrate formed clusters around the glomeruli, surrounding the renal tubules, and within the tubular epithelium (Figure 5B).

It is important to note that changes in the lymphocytic and mononuclear components may be associated with the deposition of these cells in the target organs of the reticuloendothelial system.

It appears that the G33 peptide, while stimulating a specific immune response, led to a pathological reaction in mice at the applied dosages. This reaction may be attributed to the partial homology of the amino acid sequence of the G33 peptide with mammalian proteins, particularly with mouse IFIT-2 (refer to Figure 6). The tendency of the GP protein fragment to form fibrils can result in the recruitment of this cellular factor and lead to the induction of its fibrillogenesis.

**Figure 6.**
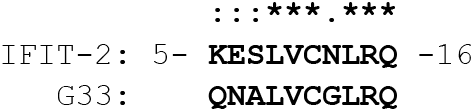
Region homologous to the G33 peptide in the mouse IFIT-2 protein (from residues 6 to 15), IFIT2_MOUSE [10]. **‘:’** denotes residues that coincide in polarity, **‘*’** denotes identical residues.

Considering the involvement of this protein in the formation of an antiviral immune response, both this protein and its orthologues can be targets for the induction of a conformational transition if GP2 or its fragments have one *in vivo*, as well as for the formation of specific antibodies that are cross-reactive with host proteins [11].

## Conclusion

It is evident that peptides in fibrillar form are capable of inducing an immune response to the full-length protein without being toxic to cells and without significantly altering the hematological analysis.

The potential of the studied fragment of the Ebola virus glycoprotein to oligomerize and form amyloid-like fibrils and intermediates could be linked to the mechanisms of action of the virus on cells, including those of the immune system. Moreover, if the native GP2 epitopes are preserved in the G33 peptide fibrils, the G33 peptide aggregates may be suitable for immunization, as fibrillogenic polypeptides are recognized to possess autoadjuvant properties [12]. However, it’s important to note that the G33 peptide subsequence is not included in the list of immunogenic pentapeptides unique to the Ebola virus that have no homology with human proteins [13], suggesting the potential cross-reactivity of a vaccine based on G33 epitopes.

## Author contributions

**Y.Z**.: Formal analysis, Investigation, Writing - Original Draft, Writing - Review & Editing, Visualization; **K.S**.: Formal analysis, Investigation, Writing - Original Draft; **M.S**.: Investigation, Writing - Original Draft; **A.A**., **E.K**., **A.T**., **M.E**., **E.E**.: Investigation; **V.E**.: Conceptualization, Investigation, Writing - Original Draft, Writing - Review & Editing, Supervision.

## Acknowledgments

This research was funded by the Ministry of Science and Higher Education of the Russian Federation (state task No. FSEG-2023-0014).

## Notes

### Competing Interest Statement

The authors have declared no competing interest.

